# Dispersal and gene flow in anadromous salmonids: a systematic review

**DOI:** 10.1101/2024.02.15.580427

**Authors:** Amaïa Lamarins, Stephanie M. Carlson, Mathieu Buoro

## Abstract

Dispersal is a ubiquitous ecological process that has been extensively studied in many plants and animals. Anadromous salmonids are an interesting system for examining dispersal, in part because of their well-known philopatric behavior, but also because of the conservation challenges related to the dispersal of hatchery-origin fish. Building on earlier work, we provide an updated systematic review of dispersal and gene flow in anadromous salmonids. In particular, we compared studies on dispersal of anadromous salmonids from wild and hatchery origins, including studies providing estimates of dispersal rates, observations of dispersal, and results from modelling studies. We reviewed 228 studies and found these were unevenly distributed among species, with Atlantic salmon, Chinook salmon, and sea trout being well-represented. Our results showcase considerable variability in estimated dispersal rates within and across studies, which is likely related to the different methodologies, dispersal propensities across species and populations, and spatial extents considered. Overall, our results confirmed a higher tendency of hatchery fish to disperse relative to wild fish, but we also found some variation across species that warrants further study. Moreover, we found that dispersal propensity tended to decline exponentially with distance, and that the drivers of dispersal varied considerably among studies. Additionally, we highlight various facets of dispersal captured across this suite of studies, including variation in terminology, methods and metrics for characterizing dispersal, and the spatio-temporal scales considered. Finally, our review revealed that few studies considered, and even fewer assessed, the implications of dispersal for the conservation and management of anadromous salmonids.

## Introduction

Dispersal is the movement of individuals between populations with potential consequences for gene flow (Ronce, 2007) and is a ubiquitous ecological process. It has been extensively studied in numerous organisms (Clobert et al., 2012; Bonte and Dahirel, 2017), including a focus on its causes and consequences (Fronhofer et al., 2023). Indeed, the proximate and ultimate causes of dispersal have been largely described, with dispersal being selected as it may reduce competition, kin interactions, inbreeding, and buffer against habitat stochasticity (Bowler and Benton, 2005). Ultimately, dispersal has several consequences for the eco-evolutionary dynamics of populations. It can limit the risk of extinction and aid in colonization, but it can also influence local population dynamics, the stability of a set of interacting populations, and adaptive potential. Conversely, effective dispersal (i.e., gene flow) can alter local adaptation and homogenise populations (Garant et al., 2007; Benton and Bowler, 2012; Webster et al., 2017; Lamarins et al., 2022). To better understand the dynamics of a local population, it is then necessary to consider its metapopulation context.

Anadromous salmonids (e.g., salmon and some trout) migrate between freshwater breeding grounds and ocean feeding grounds and are well-known for their philopatric behaviour, commonly referred to as ’homing’ (Salmenkova, 2017). Despite long distance migrations, anadromous salmonids have a remarkable ability to return to their natal river to breed. As a result, salmonids often exhibit significant local adaptation and genetic differentiation among populations (Fraser et al., 2011). However, dispersal among populations is not uncommon in salmonids. For example, using otolith microchemical analysis, large immigration rates have been measured by Donohoe et al. (2021) and Mikheev et al. (2021) on steelhead and sea trout, respectively. Perrier et al. (2010) showed evidence of long distance dispersal and recolonisation of the Seine River by Atlantic salmon. Another example comes from Leunda et al. (2013) who reported a case of long-distance dispersal of Atlantic salmon originating from the Tamar River in SW England, and captured in the Bidasoa River (NW Spain), more than 1000 km away.

Nevertheless, empirical observations of dispersal in wild populations of salmonids remain rare, possibly due to assumptions of philopatry but likely also related to the difficulty of assessing dispersal in the wild given the complex and relatively long life cycles of many anadromous salmonids. On the other hand, because of extensive habitat transformation and loss, many salmonid populations are supplemented by production in hatcheries, and the dispersal of hatchery fish into wild populations is a conservation concern. Indeed, the dispersal of hatchery-origin salmonids has received much attention because of their perceived higher dispersal propensity (Quinn, 1993) and potential impact on wild populations (Naish et al., 2007; McMillan et al., 2023). Dispersers are commonly referred to as ”strayers” in salmonids because some fish stray to explore suitable habitats or because of failure to imprinting, although this term has a maladaptive connotation impeding the study of dispersal as an adaptive life history strategy (Schtickzelle and Quinn, 2007). In short, salmonids are an interesting system to explore dispersal patterns and their implications.

Several prior reviews on the dispersal of salmonids have been published, each with a particular focus (Suppl. Mat. Table S1). First, Quinn (1993) compared dispersal of wild populations to hatchery populations, while Altukhov and Salmenkova (1994) reviewed dispersal intensity and genetic differentiation in several salmonid species. Then, Rieman and Dunham (2000) wrote an opinion paper on metapopulation dynamics in inland salmonids, while Schtickzelle and Quinn (2007) extended this perspective to include anadromous salmonids, including a discussion of the implications of considering metapopulation structure for salmon conservation. Hendry et al. (2004) provided an evolutionary perspective on homing and dispersal among salmonids. More recently, Keefer and Caudill (2014) reviewed dispersal mechanisms and rates across several species, while Bett et al. (2017) reviewed the main factors potentially influencing dispersal rates. Each of these reviews has made a significant contribution to our understanding of dispersal in salmonids.

We build on these prior studies using a systematic review to revisit the literature accumulating on dispersal in anadromous salmonids. We include facultative anadromous species such as sea trout (*Salmo trutta*) and Steelhead trout (*Oncorhynchus mykiss*), and studies focusing on both wild- and hatchery-origin fish given prior work emphasizing that dispersal may be higher in hatchery fish than in wild fish (Quinn, 1993). Our primary objective is to compare dispersal between wild- and hatchery-origin fish, and among species. In particular, we compare the dispersal estimates (emigration rate, immigration rate, and gene flow) between wild and hatchery sources, as well as the methods and spatio-temporal scales considered. In addition to empirical estimates of dispersal rates, we review the factors identified as potential drivers of dispersal, building on the work of Bett et al. (2017), and including the results of modelling studies focused on salmonid dispersal (either statistical or simulation models such as Bowlby and Gibson, 2020) because they can enhance our understanding of the dispersal process and its consequences. Finally, we discuss the implications of dispersal for the conservation and management of anadromous salmonids.

### Method for the literature review

To synthesize results from studies of dispersal focused on anadromous salmonids, we conducted a literature review on the Web of science (WOS) advanced research query using the following keywords: Species (see below) *AND* (dispers* *OR* stray* *OR* ”gene flow” *OR* ”metapopulation” *OR* ”homing”) *NOT* ”lice”. We limited this literature research from January 1, 1955, to January 24, 2023, and to articles reported in WOS, including grey literature (e.g., technical reports).

The search focused on Atlantic salmon (*Salmo salar*), Coho salmon (*Oncorhynchus kisutch*), Chum salmon (*Oncorhynchus keta*), Pink salmon (*Oncorhynchus gorbuscha*), Sockeye salmon (*Oncorhynchus nerka*), Chinook salmon (*Oncorhynchus tshawytscha*), Steelhead trout (*Oncorhynchus mykiss*), and Brown / Sea trout (*Salmo trutta*). We focused on these species because they all have at least one anadromous life history form. Because it appeared in several queries, it is important to note that individual multi-species studies were counted as several studies (i.e., one per species).

After excluding studies focused on dispersal at the juvenile life stage, resident life history forms, landlocked populations, reviews, and studies focusing on other topics or species (e.g., parasites, introgression, physiology), we retained 334 studies that focused on dispersal in salmonid populations that were at least partly anadromous. Next, we excluded studies (n=122) that were solely concerned with the evolutionary history and genetic differentiation between populations, without providing estimates of dispersal, bringing our total to 212 studies. Finally, we added 16 references to the database using a snowball approach, in which we identified relevant studies from studies identified from our WOS search. Throughout, we retained only those studies cited in previous reviews or original studies that estimated or suggested dispersal in wild or hatchery populations. Following these filtering steps, we retained 228 studies for inclusion in our review (Supplementary Material 1, online database https://doi.org/10.57745/RCMAGH).

We classified the 228 retained studies according to the source of dispersal (wild- or hatchery-origin), study approach (estimated one or more dispersal metrics, observations, modeling), and species (see section below, Figure 1). From the studies providing estimates of dispersal (n=183), we collected the estimated rates and classified them as one of three metrics of dispersal: *emigration rate* (or ”donor population stray rate” *sensu* Keefer and Caudill, 2014), *immigration rate* (or ”recipient population stray rate” *sensu* Keefer and Caudill, 2014), and *gene flow* (or ”effective migration”). Regarding the studies focused on gene flow, we did not analyze the estimates of the number of migrants per generation because they were not comparable between studies; rather, only migration rates were considered. For each study, we identified the approaches and methods (e.g., tagging, micro-chemistry) used, the spatial and temporal scales of measurements, and reported the distance between donor and recipient populations (including hatcheries and natural spawning areas). If a distance measurement was not specified in the study, we measured the shortest waterway distance, which is the distance separating two populations along the coastline or the river system (Supplementary Material 2). We also reported whether studies investigated the potential drivers of dispersal, including individual (e.g., sex, age), population factors (e.g., density), and environmental factors (e.g., water temperature, discharge). From the 228 retained studies, we also extracted (and reported) the frequency of different terms appearing in each manuscript referring to aspects of dispersal to explore the language used to characterize dispersal in studies focused on wild- and hatchery-origin fish. Finally, we determined whether the implications of dispersal for conservation and management were discussed or investigated.

**Figure 1:**
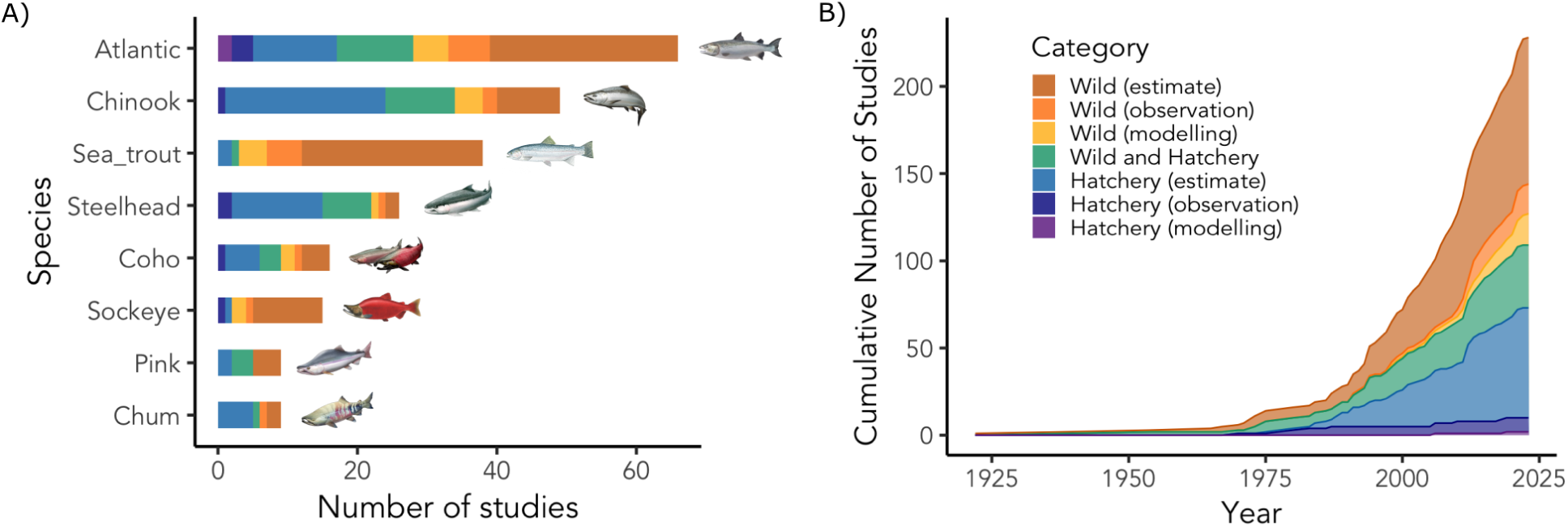
A) Number of studies of each category per species. B) Cumulative number of studies over time per category (merged across all species). © Illustrations: 2022 California Trout Inc.

## Query overview

### Species representation

Across the 228 studies included in our review, we found that species were not evenly represented (Figure 1 A). In particular, Atlantic salmon was the most common species that emerged through our systematic review of dispersal in anadromous salmonids (29% of studies), followed by Chinook salmon (21.5%), sea trout (16.7%), steelhead trout (11.4%), Coho salmon (7%), Sockeye salmon (6.6%), and, finally, Pink and Chum salmon (3.9% each). Keefer and Caudill (2014) also reported a higher number of studies focusing on the dispersal of Atlantic salmon, Steelhead, Chinook salmon, as compared to Coho, Sockeye, Pink, and Chum salmon. It is noteworthy that Pacific salmon, and particularly Pink salmon - a highly dispersive species and currently invasive in Europe (Sandlund et al., 2019) - were underrepresented in the studies we encountered.

### Terminology

The diversity of terms used by studies to describe dispersal among populations illustrates the many facets of dispersal. In examining studies of dispersal from wild origin, we found that the most common terms used to describe aspects of dispersal within manuscripts included *migration* (-migra-, 40% of studies), *straying* (stray-, 34%), *gene flow* (10%), *dispersal* (dispers-, 9%), and *metapopulation* (metapop-, 4%). In contrast, studies of hatchery-origin dispersal predominantly favored the term *straying*, used by 70% of studies, while *migration* was the primary term in 28% of studies. The use of *dispersal* and *gene flow* was notably less frequent in hatchery-origin dispersal studies, appearing in only 2% and 1% of cases, respectively.

The term *migration* was frequently used by genetic studies to refer to measures of migration rate (gene flow), immigration rates, or number of migrants. However, this term is not restricted to dispersal behavior as it could also refer to migration within river or migration at sea. *Gene flow* might be more appropriate but was less predominant than *migration* in our review. *Straying* was also prevalent in our review and especially used by studies focusing on hatchery fish. *Dispersal* might be used interchangeably with straying but it was less predominant. However, dispersal (or straying) and gene flow (or migration) cannot be used interchangeably as gene flow is not a proxy for dispersal. In fact, it may be lower or higher than the dispersal rate due to reduced or enhanced reproductive success and fitness of immigrants. Consequently, in this manuscript, we used the term ”philopatry” instead of ”homing,” ”dispersal” instead of ”straying,” and ”gene flow” or ”effective dispersal” instead of ”migration”.

### Classification of studies by source: wild- vs. hatchery-origin

Our review revealed that empirical studies focusing solely on the dispersal of wild-origin anadromous salmonids represented 44% of the retained studies, most of them providing estimates of one or more dispersal metrics (84 studies, Figure 1 ”Wild (estimate)”). Studies providing evidence of dispersal without estimating one of the dispersal metrics (17 studies, Figure 1 ”Wild (observation)”) were generally based on relatively small sample sizes and occasional observations, but provided important information by adding further evidence to dispersal in wild populations, including one-off observations of long-distance dispersal (Perrier et al., 2010; Leunda et al., 2013). We identified a few modelling studies (statistical or simulation models) - focused on dispersal of wild populations of salmonids (8%, 18 studies, Figure 1 ”Wild (modelling)”), mainly published since 2010. Most modelling studies have focused on Atlantic salmon, brown trout, and Chinook salmon, with a smaller number focused on Coho salmon, Sockeye salmon, and Steelhead trout. We encountered no models in our systematic review that focused on Pink salmon or Chum salmon.

In contrast to the dispersal of wild-origin fish, monitoring the dispersal of hatchery-origin fish is widespread because of management and conservation implications, and is often facilitated by large-scale tagging programs focused on hatchery fish. Hatchery tagging programs may thus provide insights into the causes and consequences of dispersal, although it is important to recognize that hatchery fish may or may not be good surrogates for wild fish. Overall, a significant proportion of studies (32%) focused on the dispersal of hatchery-origin fish including farm escapees (n=4), or transplanted fish. In particular, 63 studies estimated one or more of the dispersal metrics, while 8 studies provided observations only, and 2 modelling studies evaluated the impact of hatchery dispersal on wild populations (Figure 1 ”Hatchery (estimate)”, ”Hatchery (observation)” and ”Hatchery (modelling)”). Most of these studies focused on (in order of their representation in our review pool): Chinook salmon, Steelhead trout, Chum, Coho, Pink, and Atlantic salmon. The focus on characterizing dispersal of hatchery fish has been driven by hatchery management’s objectives, including understanding efficiencies (or ”loss” of fish in the stocking program), concerns about fulfilling hatchery broodstock, and/or concerns about dispersal to wild populations. Earlier reviews of dispersal highlighted the higher dispersal propensity of hatchery- compared to wild-origin fish (Quinn, 1993; Keefer and Caudill, 2014), and our review shows that the interest in dispersal of hatchery vs. wild fish persists (Figure 1 B).

Importantly, 36 studies (16%, Figure 1 ”Wild and Hatchery”) provided estimates of dispersal from both wild- and hatchery-origin fish, allowing for a direct comparison of dispersal in hatchery and wild fish of the same species, using the same methods and dispersal metrics, and for a similar temporal and spatial context.

### Metrics of dispersal

In this review, we focused on the three metrics of dispersal together: emigration, immigration and gene flow. While emigration rates are often measured both from wild and hatchery populations, information on emigration alone does not reveal the full impact of dispersers. For example, even a small emigration rate from a large population can have a substantial impact on the demographics of a neighbouring smaller population (Keefer and Caudill, 2014) - e.g., when large proportions of hatchery-origin fish disperse into wild populations (Knudsen et al., 2021). Therefore, immigration rates may be more relevant for evaluating the consequences of dispersal (Bett et al., 2017), particularly in the context of salmon metapopulations which can include a mixture of hatchery supported and wild populations. However, immigration rates do not inform on the reproductive success of immigrants and the genetic consequences of dispersal. Consequently, measures of gene flow are also complementary although they provide only a partial picture of the impacts of contemporary dispersal.

Our review revealed that most studies of wild-origin fish measured emigration rates from wild populations (39%) and gene flow (33%), closely followed by the immigration rate into wild populations (28% of studies, Figure 2). Keefer and Caudill (2014) found that studies focused disproportionately on emigration rates, and our results suggest that this is changing (with better balance between studies focused on emigration from donor populations and immigration into receiving populations). However, when we add information on source (hatchery or wild), we found that studies based on hatchery-origin fish mainly estimated emigration rates (70%), much more similar to the earlier results of Keefer and Caudill (2014), with a smaller proportion estimating immigration rates (26%) and even fewer focusing on gene flow (4%). The limited number of studies estimating gene flow between hatchery and wild populations may seem surprising, but it may reflect our choice to not retain studies of genetic introgression.

**Figure 2:**
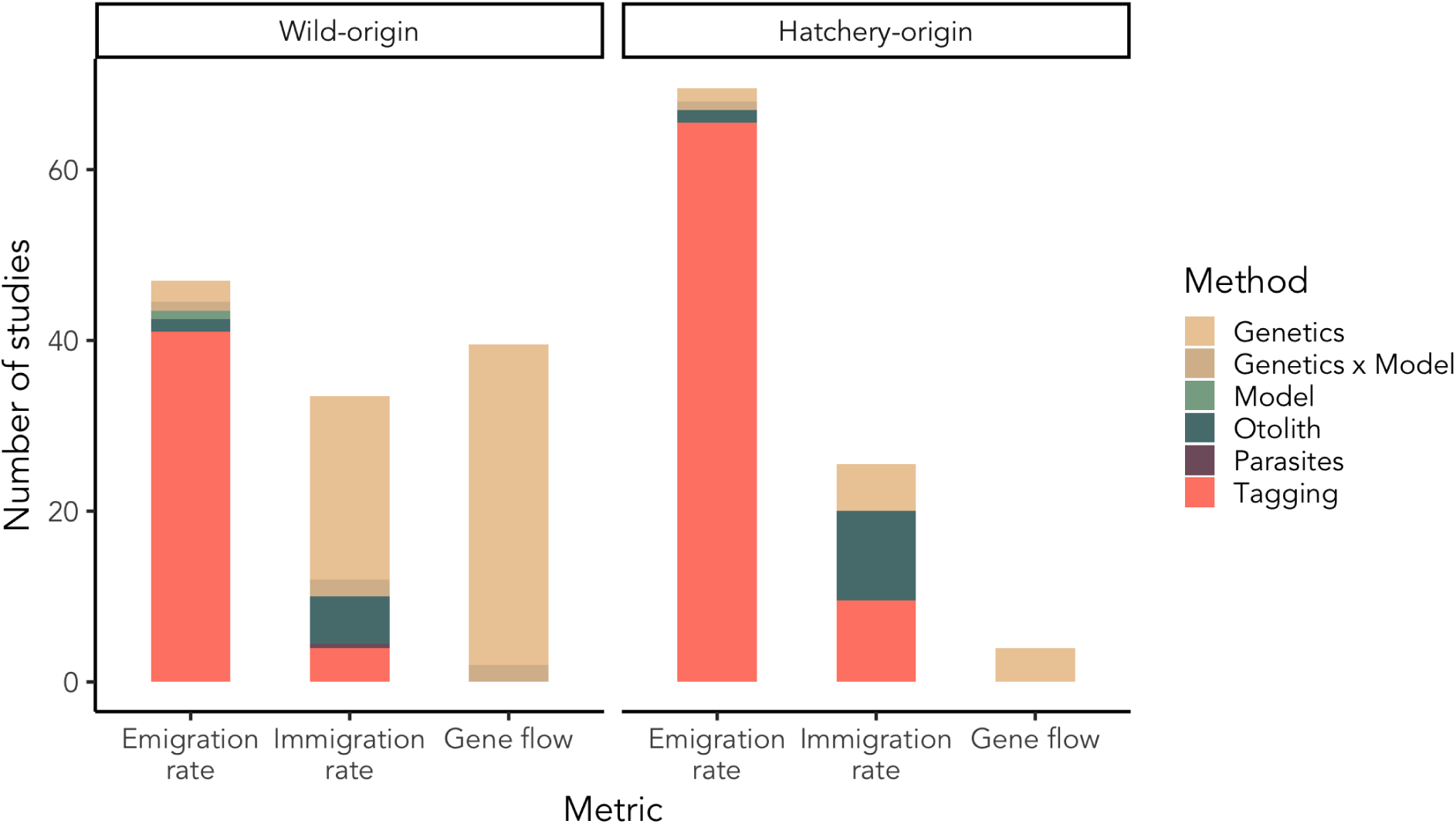
Number of studies providing estimates of emigration rate, immigration rate, and gene flow, broken down by method (e.g., genetic, tagging) and presented separately for each source (wild-origin vs. hatchery-origin).

Not surprisingly, using a combination of approaches and metrics holds promise for overcoming the limits of any given approach or metric and providing a more nuanced understanding of the role of dispersal within a given salmon system. However, in our review, we found that only 14% and 9% of studies of dispersal of wild- and hatchery-origin, respectively, used two or more metrics (e.g., Tallman and Healey, 1994; Mortensen et al., 2002; Masson et al., 2018).

### Dispersal rates: wild- vs. hatchery-origin

A primary goal of our study was to compare dispersal rates between wild- and hatchery-origin fish. To do so, we compared the distribution of dispersal rate estimates between wild- and hatchery-origin fish gathered from 183 studies (Figure 3), although it is important to recognize that the estimated rates of dispersal were measured in different ways and in various contexts (e.g., spatial and temporal) and should be interpreted with caution. Based on these data, we found that the emigration rate from hatchery populations (median: 7%, mean: 15.4%, *sd*: 21.2%) was significantly higher than from wild populations (median: 3.7%, mean: 8.3%, *sd*: 12.7%; Kolmogorov-Smirnov test: D = 0.18, p <0.001). This result was expected based on the results of prior research (e.g., Keefer and Caudill, 2014; Pearsons and O’Connor, 2024). However, we found that this pattern did not hold true across all species (Figure 3 A). Rather, the emigration rate was higher for hatchery populations than for wild populations of Atlantic salmon, Chinook salmon, and Steelhead trout, but the opposite was true for Pink, Coho, Chum salmon, and sea trout, although we should note that there were few values reported from wild populations for some of these latter species (especially Chum salmon [n=4] and sea trout [n=7]). A subset of studies provided a direct comparison between estimated rates of emigration for hatchery- and wild-origin fish from similar spatial and temporal contexts. For example, Jonsson et al. (1991, 1994, 2003) and Hansen et al. (1993) showed from 1.4 to 10 times higher emigration rates from hatchery Atlantic salmon, compared to wild fish from the river Imsa. Jonsson and Jonsson (2014) reported higher emigration rates in hatchery sea trout (7 times higher than wild), while Labelle (1992) reported that emigration rates for hatchery Coho salmon were three times higher compared to wild fish. A parentage analysis of Chinook salmon led by Ford et al. (2015) estimated an emigration rate of 29% from the Chiwawa Hatchery compared to only 4.1% for the wild population. In contrast, Keefer et al. (2008b) did not observe higher emigration from hatchery Chinook salmon released in river compared to wild Chinook salmon within the Columbia River Basin, although they did find higher emigration of hatchery Steelhead trout relative to wild fish. The results of Thedinga et al. (2000) on Pink salmon also did not support higher emigration rates from hatchery fish compared to wild fish in southeastern Alaska.

**Figure 3:**
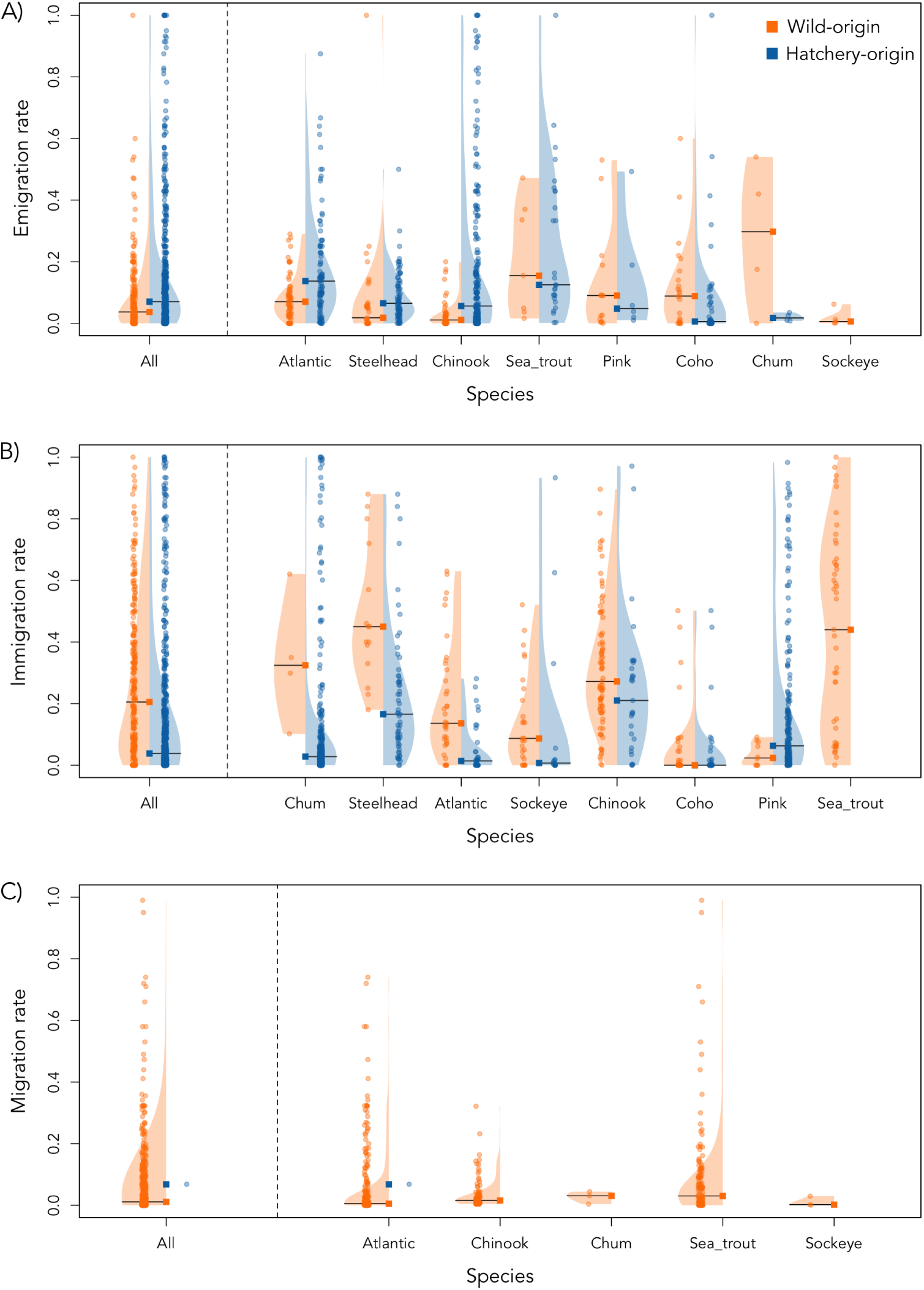
Distributions of estimated A) emigration, B) immigration, and C) migration rates among species, grouped by the source of fish (wild- or hatchery-origin). Migration rates exclude estimates of number of migrants per generation (*Nm*). Each point is an estimated value and the horizontal lines and squares show the median value.

Conversely, the overall immigration rate of hatchery-origin fish, measured in both wild and hatchery recipient populations, was significantly lower (median: 3.8%, mean: 13.7%, *sd*: 22.5%) than that from wild-origin fish (median: 20.5%, mean: 26.6%, *sd*: 24.9%; Kolmogorov-Smirnov test: D = 0.34, p <0.001), with the exception of Coho and Pink salmon (Figure 3 B). This result is surprising given the higher tendency of hatchery fish to disperse and considering that hatcheries are typically located in impacted watersheds with reduced levels of natural production. This result may be explained by a bias in the immigration rates reported, which often included wild and small populations, which are potentially easier to monitor and subject to higher immigration rates from wild-origin fish. Keefer and Caudill (2014) also highlighted the challenges inherent in investigating immigration rates and thus making valid comparisons. In our review, a very few studies provide a direct comparison of the immigration rate from wild- and hatchery-origin salmonids within a same system. In the Cedar River, after several years of natural recolonization of a newly accessible habitat, Anderson et al. (2015) measured similar proportion of hatchery-origin and wild-origin Chinook salmon immigrants (about 1/3), while in Coho salmon, most of the immigrants, which represented about a third of the population, were wild-origin immigrants. The natal origins of Chinook salmon reestablishing a population in a restored small creek from California’s Central Valley were identified in Willmes et al. (2021) and overwhelmingly attributed to Central Valley hatcheries (88%).

Unfortunately, it was not possible to compare effective migration rates between wild- and hatchery-origin fish (Figure 3 C), because gene flow has not been measured in all species, and only one study measured gene flow from hatchery fish (m=6.8%; Vasemägi et al., 2005). However, it is interesting to note that migration rates estimated from genetic data (median: 1.1%, mean: 5.66%, *sd*: 11.8%) were lower than immigration rates from wild-origin fish. This suggests that immigrants of wild-origin (and/or their descendants) could have a lower fitness (reproductive success and/or offspring survival) than local fish, which in turn results in a low level of gene flow (Peterson et al., 2014). For example, Tallman and Healey (1994) found low levels of gene flow in Chum salmon despite both high estimated rates of emigration and immigration, suggesting low reproductive success of immigrants. Similarly, despite an immigration rate of 12.5% in a population of Atlantic salmon, Mobley et al. (2019) conducted a parentage analysis and showed only 3.7% of juveniles were assigned to dispersing adults, again hinting at low reproductive success of immigrants.

In conclusion, while our global analysis indicates a higher tendency for hatchery fish to disperse compared to wild fish, the patterns appear to be species- and context-dependent. It is important to note that the average emigration rate may appear low, suggesting that dispersal in salmonids is negligible. However, this average combines measurements from various contexts (e.g., distances), even though dispersal can be highly variable and significant among nearby populations and within populations across years under different environmental conditions.

### Drivers of dispersal

Overall, estimates of emigration, immigration, and migration rates spanned the full range between 0% and 100%, and showed a high degree of variability within and among species (Figure 3).

### Species propensity to disperse

Variation in estimated rates among species could reflect different dispersal propensities. However, across all three metrics (emigration rate, immigration rate, migration rate), we found no clear pattern among species (Figure 3 A-C). However, this result may be a consequence of comparing studies from different systems (and spatial networks) and using different methods. A few studies (n=11) provided direct comparisons of dispersal rates between different species from the same river system or the same hatchery. For example, Shapovalov and Taft (1954) showed higher emigration rates in Coho salmon compared to steelhead trout in both wild- and hatchery-origin fish, while similar rates have been measured between wild Chum and Chinook salmon by Lin et al. (2011). Keefer et al. (2005, 2008b) reported higher rates of emigration for steelhead trout compared to Chinook salmon from both wild- and hatchery-origins in the Columbia River Basin, USA, while Pearsons and O’Connor (2020) did not find differences in the average rate of emigration between these two species in the upper part of the same watershed. On the contrary, Westley et al. (2013) measured higher emigration rates from Chinook salmon compared to Steelhead and Coho coming from similar hatcheries in the Columbia River Basin, USA in the same years. Surprisingly, Ebel et al. (1973) and Slatick et al. (1975) found no evidence of dispersal from both wild and hatchery Steelhead and Chinook salmon. However, this result may be explained by the non-exhaustive monitoring conducted. Regarding immigration rates, Anderson et al. (2015) found similar proportion of wild-origin immigrants into populations of Coho and Chinook salmon. Knudsen et al. (2021) and Brenner et al. (2012) measured higher proportion of hatchery fish into Pink salmon populations compared to Chum salmon populations, which were even higher than for Sockeye salmon. So overall, even studies providing direct comparisons among species report variable results, sometimes finding significant differences between species, and in other cases finding little variation between species. As studies of dispersal in anadromous salmonids accumulate, this should allow an updated exploration of patterns and the drivers of differences among species, both within and across basins.

### Relationship with distance

Dispersal is a behavior associated with several potential costs (Bonte et al., 2012), including the risk of failing to identify suitable habitats or partners. These risks may be heightened with increasing dispersal distances. As a consequence, the fitness of dispersers is assumed to be low if the recipient population is far from the natal river (i.e., isolation by distance), or if the recipient population is phenotypically and/or genetically differentiated (i.e., isolation by adaptation). It is thus commonly assumed that dispersal rates decrease as distance between populations increases. The relationship is commonly depicted using a dispersal kernel that considers euclidean distance as the primary factor (Nathan et al., 2012). In our review, we identified a few studies that illustrated the dispersal kernel by providing the distribution of dispersal distances among emigrants, such as Berg and Berg, 1987 (wild sea trout), Jonsson et al., 2003 (wild Atlantic salmon), Thedinga et al., 2000 (wild Pink salmon), Jonsson and Jonsson, 2014 (wild and hatchery sea trout), and Unwin and Quinn, 1993 (hatchery Chinook salmon). The results from these studies are presented in Figure 4 and show a significant pattern of exponential decline in distance moved among fish that disperse, including data from studies of both wild- and hatchery-origin fish (Suppl. Mat. Table S2). However, long distance dispersal (up to 2000km, e.g., Hansen, 2006) has been documented in several studies and a reverse dispersal kernel has also been suggested in Hansen (2006), based on releases of farmed Atlantic salmon from Norwegian sea cages, meaning a larger proportion of dispersers moving long distances.

**Figure 4:**
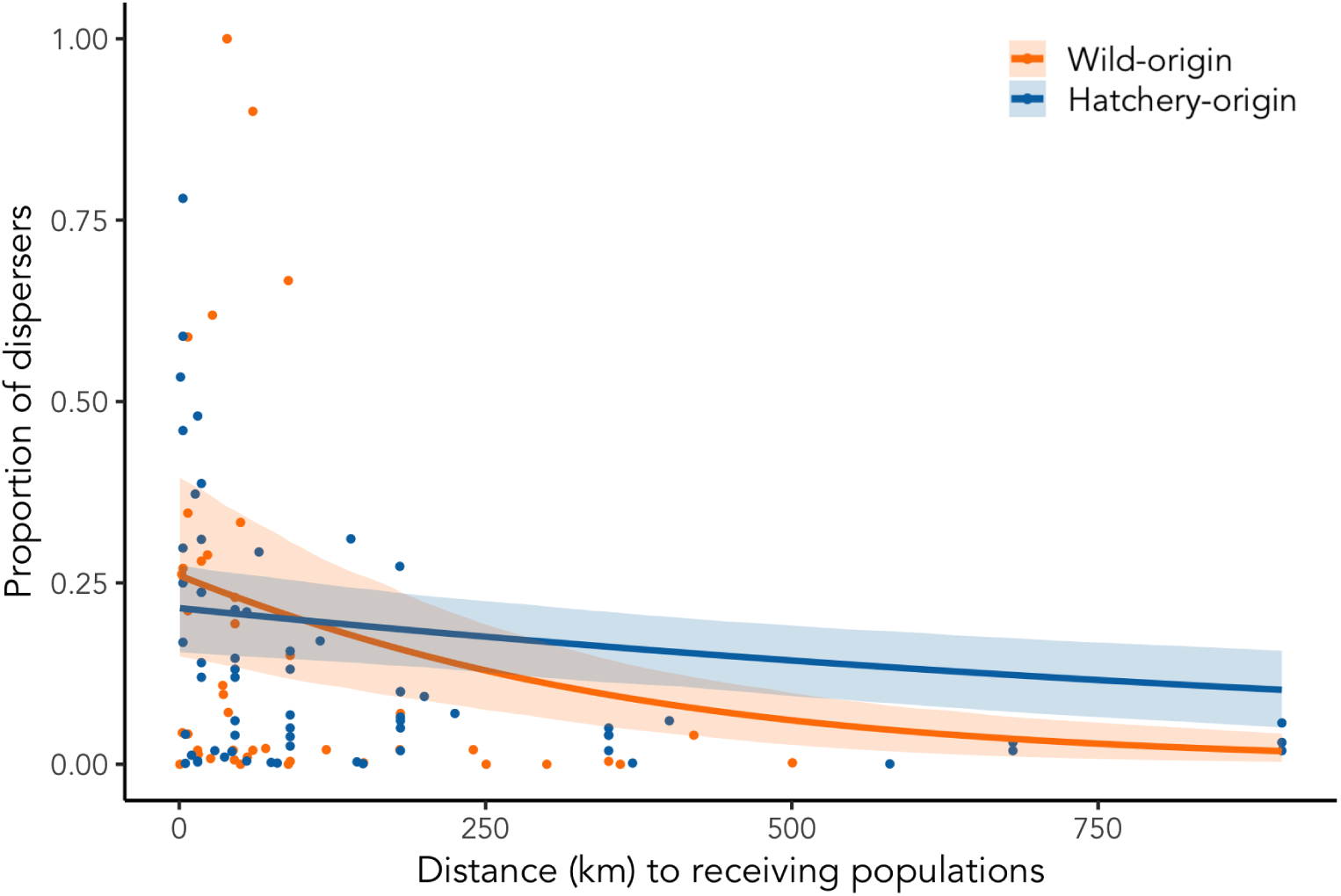
For the subset of dispersers, the proportion of dispersers as function of the shortest waterway distance travelled to their receiving populations, by source (wild- and -hatchery origin). Fitted curves correspond to beta regressions with logistic link (Suppl. Mat. Table S2).

We also attempted to explore the relationship between the three dispersal metrics (i.e., emigration, immigration, and migration rates) and distance (averaged distance for emigrants and immigrants travelled from source to final destination) reported across all studies and species (Suppl. Mat. Figure S1). The relationship between emigration rates and distance to receiving populations was not significant, and this result was true for both studies based on wild fish as well as studies based on hatchery fish (Suppl. Mat. Table S2). Nevertheless, we did note a trend for a decrease in emigration strength with increasing distance, with the highest emigration rates associated with the smallest distances, which might indicate spatially structured population dynamics. Moreover, we noted the distribution of dispersal distances included higher values for hatchery fish as compared to wild fish. However, inspection of the immigration patterns revealed higher long distance dispersal (up to 710 kms) in wild fish compared to hatchery fish, and significant - but opposite - relationships with distance from source populations between wild (significantly positive) and hatchery (significantly negative, Suppl. Mat. Table S2). However, it is important to interpret these results with caution because certain systems played a large role in driving these patterns. For example, the high wild-origin proportion of immigrants from long distance populations were mostly reported from large river systems in North America, e.g., the Fraser and Columbia rivers (Hess and Matala, 2014; Bett et al., 2017; Matala et al., 2017). Gene flow, which results from immigration and the reproductive success of immigrants, also slightly decreased with increasing distance among wild populations although the relationship was not significant (Supp. Mat. Table S2). Indeed, gene flow is likely to be reduced with increased distance between populations because of local adaptation which may limit the reproductive success of dispersers.

Overall, our review highlights that distance is a simple and convenient descriptor of dispersal patterns. However, distance does not capture the underlying mechanisms of the emigration process, nor the choice of the recipient population or the reproductive success of immigrants. For example, immigration rates may be more likely influenced by the relative population size than by distance. If a small population receives immigrants from a large donor population, the immigration rate might be high despite a large distance separating the populations in question. Moreover, the attractivity of populations may also influence this pattern (i.e., collective behaviour, Berdahl et al., 2016), with large populations attracting relatively more immigrants than small populations. Together, this suggests a need for further studies of additional putative drivers involved in dispersal for an improved understanding of its causes and consequences. Especially, individual attributes (e.g., sex, age), population (e.g., density) or environmental factors (e.g., water flow) may affect emigration and immigration patterns via several processes, which we describe below.

### Individual, population and environmental drivers

Few studies have examined the primary factors that influence dispersal propensity in salmonids, which limits our understanding of the drivers of dispersal (see Keefer and Caudill, 2014; Bett et al., 2017 for reviews). Here, we quantified the studies that assessed the influence of one or more factors on dispersal propensity of wild- and hatchery-origin fish separately. Only a small proportion (19% overall) of studies estimating wild-origin dispersal rates evaluated factors promoting dispersal, such as individual characteristics (e.g., sex), environmental conditions (e.g., temperature), or population density (Figure 5). However, relatively more studies (53%) on hatcheries evaluated potential drivers of dispersal, possibly reflecting the relative ease of tagging hatchery fish prior to release and challenges of assessing dispersal in wild populations especially over large areas and through time.

**Figure 5:**
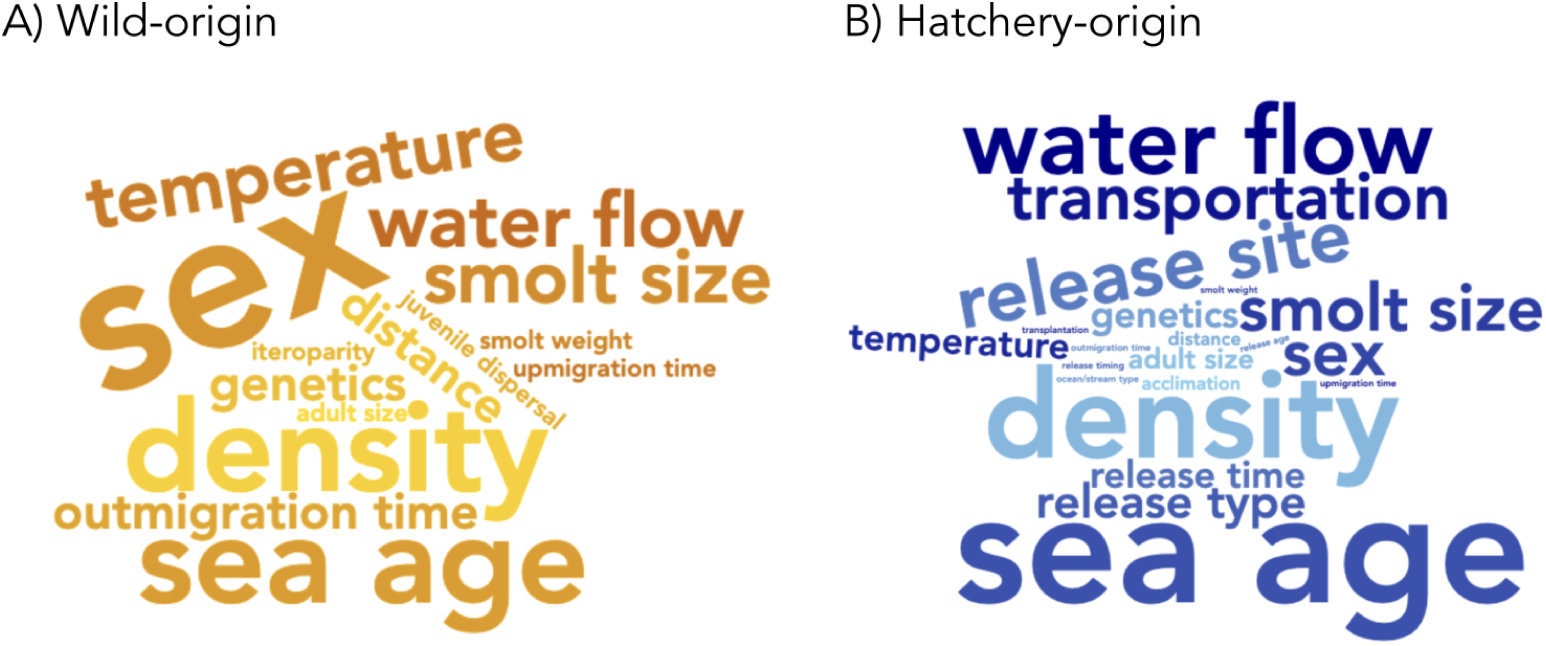
Wordcloud of the factors identified as potential drivers of dispersal (emigration or immigration patterns), their size being proportional to their occurrence among studies, presented separately for A) wild- and B) hatchery-origin fish.

Our review revealed that the primary factors that were examined as potential drivers of dispersal, in both wild- and hatchery-origin fish, were individual sex, body size, and sea age (Figure 5). Because theoretical studies suggest that sex-biased dispersal is widespread and contingent on mating systems and kin competition (Trochet et al., 2016; Li and Kokko, 2019), we explored evidence for sex-biased dispersal. Overall, we found mixed evidence of sex-biased dispersal, with some studies reporting no effect of sex (Thedinga et al., 2000; Palstra et al., 2007; Consuegra and Garía de Leániz, 2007; Ford et al., 2015), whereas others provided results consistent with male-biased dispersal, both in wild populations (Bekkevold et al., 2004; Hamann and Kennedy, 2012) and in hatchery populations (Hard and Heard, 1999), and others consistent with female-biased dispersal (Pollock et al., 2020). Body size may also affect dispersal due to potential size-dependent costs of dispersal (e.g., due to differences in the arduousness of migration), while sexual selection can lead to differences in body size among populations. No difference was found between the length of hatchery sea trout dispersing and returning to their home river in Degerman et al. (2012), whereas Pollock et al. (2020) showed that larger hatchery Chinook salmon were more likely to be within-river dispersers than smaller fish. However, size at downstream migration (i.e., smolt size) was found to be negatively correlated with Coho salmon and Steelhead trout emigration rate (Shapovalov and Taft, 1954) but this relationship was positive for hatchery reared Steelhead trout in Clarke et al. (2014) and not significant in hatchery sea trout and Chinook salmon (Jonsson and Jonsson, 2014; Unwin and Quinn, 1993). Finally, time spent in river or at sea can also affect the olfactory imprinting of juvenile salmon in the river and the olfactory memory of adults at sea, and therefore their ability to distinguish between water sources (Dittman and Quinn, 1996). Again, we found inconsistent patterns across studies regarding the influence of sea age on dispersal propensity. For example, while no difference across sea ages was observed in Atlantic and Chinook salmon (Potter and Russell, 1994; Ford et al., 2015), Källo et al. (2022) and Hard and Heard (1999) instead showed a negative relationship between emigration rate and time at sea in wild sea trout and hatchery Chinook salmon, whereas Jonsson et al. (2003) and Unwin and Quinn (1993) measured higher dispersal with sea age both in wild Atlantic salmon and hatchery Chinook salmon. However, it is difficult to disentangle the effects of age versus other, potentially correlated, factors. For example, females are likely to be older (spending more time at sea) and larger, suggesting it is important to consider both sex and age. More generally, studies exploring multiple factors, and their interactions, would be a valuable addition to the growing body of literature exploring drivers of dispersal.

Another important factor influencing dispersal propensity is density. For example, Jonsson et al. (2003) and Hard and Heard (1999) provided evidence of a negative relationship between returning adults density and emigration rate of hatchery-origin Atlantic and Chinook salmon, supporting the hypothesis of collective behaviour presented in Westley et al. (2015) and Berdahl et al. (2016). However, the release group size of hatchery juvenile Chinook salmon induced more frequent dispersal in Sturrock et al. (2019), while juvenile rearing density did not have any effect on dispersal propensity in Clarke et al. (2013).

Regarding environmental factors, ocean and freshwater temperature has also been explored as a potential driver in 6 studies from our literature review. This body of work revealed a positive correlation between the North Atlantic ocean temperature, immigration rate, and gene flow on wild Atlantic salmon populations (Valiente et al., 2010; Horreo et al., 2011) whereas a negative correlation was described between Chinook salmon dispersal and the Pacific Decadal Oscillation climate index in Westley et al. (2015) and Sturrock et al. (2019). Inconsistent results were also found with freshwater temperature during upriver migration. While Bond et al. (2017) reported a positive correlation between river temperature and emigration rate from hatchery Chinook salmon, no relationship with air temperature was found in a study of sea trout dispersal by Degerman et al. (2012). Water flow or discharge may also influence the choice of river destination for salmon returning, with the possibility that higher discharge may be an attractant (Quinn et al., 1991). Here again, we found an inconsistent pattern in terms of the influence of discharge across studies in our review. In particular, while Labelle (1992) and Degerman et al. (2012) explained the high immigration rate of Coho and sea trout populations by the river’s large discharge, Tattam and Ruzycki (2020) found no effect of river discharge in hatchery steelhead populations. Schroeder et al. (2001), Jonsson et al. (2003), Westley et al. (2015) and Bond et al. (2017) did not find a significant relationship between discharge and emigration rate across different salmonid species.

The influence of hatchery practices on dispersal propensity has also been investigated specifically for hatchery dispersal studies (Figure 5 B), including release location (e.g., offshore vs. estuary), transport, release timing, with sometimes conflicting results. While Eriksson and Eriksson (1991), Jonsson et al. (1994), Hvidsten et al. (1994), and Bond et al. (2017) showed no influence of release location, Heggberget et al. (1991), Candy and Beacham (2000) and Jonsson and Jonsson (2021) showcased increased dispersal propensity for smolts released offshore / at the river mouth (Lasko et al., 2014). Different release strategies, in particular barging or trucking of reared smolts to downstream locations before release have also been showed to increase dispersal in many studies (Keefer et al., 2008b; Bond et al., 2017; Sturrock et al., 2019; Tattam and Ruzycki, 2020). Moreover, Pascual et al. (1995) and Sturrock et al. (2019) showed an effect of smolt release timing on dispersal, while Unwin and Quinn (1993) found that release timing influenced dispersal between basins but not within basins. Finally, despite its evidence in many other organisms (Saastamoinen et al., 2018), the genetic basis of dispersal has been poorly studied in wild populations of salmonid fishes, but a few studies with rearing and transportation treatments of different stocks suggested a heritable component of dispersal behaviour (Mclsaac and Quinn, 1988; Hard and Heard, 1999; Candy and Beacham, 2000; Jonsson and Jonsson, 2017).

### Other sources of variability in dispersal rates

Several other factors, including methodological factors, spatial and temporal context, potentially contribute to the large variation in dispersal rates observed.

### Methodological factors

The methods used to estimate dispersal rates are important sources of variability across studies. To estimate the dispersal metrics, a diversity of methods have been used (Figure 2), sometimes in combination (e.g., 9% and 5% of studies focused on dispersal of wild- and hatchery-origin fish, respectively, using a combination of methods). Genetic methods represent a large proportion of the methods used to estimate the dispersal of wild-origin fish, accounting for approximately half of the studies conducted on most species, and an even larger proportion for sea trout. They were primarily used to estimate gene flow (e.g., F statistics, Wright, 1951; allelic frequencies, Wang and Whitlock, 2003; Wilson and Rannala, 2003) and immigration rate (e.g., genetic assignment, parentage analysis). In contrast, genetic methods were rarely used in studies focused on dispersal of hatchery-origin fish (only 11%), perhaps because of the ease of tagging hatchery fish relative to wild fish and the large-scale tagging programs connected to many production hatcheries. Instead, we found that the majority of studies exploring dispersal of hatchery fish (76%) relied on tagging methods (e.g., Carlin tags, fin clips, Coded Wire Tags). Studies relying on physical tags also represented a significant proportion of the methods used to study the dispersal of wild-origin fish (37.5%). Overall, we found that physical tags have been used primarily to estimate emigration rates (Figure 2). Calcified structures such as otoliths has been used in a few studies of wild- (5.8%) and hatchery-origin (12%) fish to estimate emigration and immigration rates, through micro-chemistry (e.g., Hamann and Kennedy, 2012; Turcotte and Shrimpton, 2020) or thermal marking analysis (e.g., Mortensen et al., 2002; Brenner et al., 2012). One study also used parasite prevalence to estimate the immigration rate in Sockeye salmon (Quinn et al., 1987).

Prior work has emphasized the benefits and limitations of the different direct and indirect methods for estimating dispersal metrics provided by genetic and tagging approaches (e.g., Koenig et al., 1996; Bossart and Pashley Prowell, 1998; Bullock et al., 2006). Ideally, different methods (e.g., genetic *vs.* tagging) should be applied to the same system to estimate the same metric (e.g., immigration rate). But we only found a few studies reporting differences in immigration rates by using different methods (Bartron et al., 2004; Pedersen et al., 2007; Hess et al., 2014, genetic vs. tagging; Kuparinen et al., 2010, tagging vs. model). We found no studies estimating emigration rates using more than one method. However, modelling approaches have been used in combination with genetic methods to estimate all three metrics (emigration, immigration, and gene flow) in a few studies (Bekkevold et al., 2004; Lin et al., 2011; Perrier et al., 2013).

### Exploratory behavior

Keefer and Caudill (2014) discussed the challenge of distinguishing ”permanent” vs. ”temporary” dispersal. Indeed, we often get only a partial view of dispersal behavior, because apparent observations of dispersal or philopatry (e.g., a fish in a non-natal river, or a fish in a natal river) may just represent temporary transit / exploration. This exploratory behavior was the focus of Peterson et al. (2016), who showed that Sockeye salmon dispersers were sometimes initially observed in their natal rivers and then later moved to non-natal rivers to spawn, while philopatric individuals were rarely observed in non-natal rivers. Keefer et al. (2008a) also found that a significant proportion of Chinook salmon temporarily navigated into non-natal tributaries before returning to their natal river to breed. The results of Chat et al. (2022) revealed a high proportion of sea trout immigrants were found close to the mouths of three distinct rivers, also suggesting temporary dispersal, while Richins and Skalski (2018) quantified the relationship between the probability of dispersal and the ”overshooting” rate across both wild- and hatchery-origin steelhead trout. Given the potential for (and observations of) exploratory behavior, it is important to consider that the spatial configuration and distance to other populations may then influence both the emigration and immigration rates because isolated populations are less likely to receive or produce dispersers, or to be used temporarily as staging or refuge habitat.

### Spatial and temporal scale of dispersal

Perhaps not surprisingly, our review revealed considerable variation among studies in the spatial and temporal scales at which dispersal was measured. This has consequences for estimates of dispersal and the interpretation of these estimates, as illustrated by the strong differences in dispersal propensity reported within studies between different river systems, or even among populations from the same river system (e.g., Labelle, 1992; Hess and Matala, 2014). First, spatial dispersal of anadromous species can occur both among basins (i.e., different estuaries) and within basins (i.e., common estuary) and the spatial configuration of populations can influence dispersal behaviour and propensity. We reviewed the spatial scale of studies to determine whether the extent of dispersal among and within basins was similar. 54% and 52% of the studies of wild-and hatchery-origin fish investigated dispersal among basins, 34% and 41% within basins, and 10%-7% considered both. It is plausible that dispersal within a basin might be higher than among basins. We find some support for this idea from the overall results of emigration rates in wild populations, which are higher within basins compared to among basins, despite a low number of estimates from within basins (Suppl. Mat. Figure S2). For a single system, estimated rates can also vary strongly depending on the spatial scale considered. This is very well described by Pearsons and O’Connor (2020) who showed emigration rates of Chinook salmon and Steelhead trout were lower at the basin scale (on average 0.23% and 0.2%, respectively) compared to sub- basin scale (2.36% and 2.38%) and tributary scale (2.42% and 4.57%). Hamann and Kennedy (2012) also showed immigration rates of Chinook salmon at two locations were higher when considering a fine scale (40% and 73%) than a coarse scale (12% and 18%), while Chat et al. (2022) showed a negative relationship between immigration rate and distance to river mouth of sea trout across several river systems.

Additionally, dispersal intensity may strongly vary over time, and temporal variation in dispersal may have substantial eco-evolutionary consequences (Peniston et al., 2024). However, most of the studies (60% and 47% for wild- and hatchery-origin fish) reported rates that were pooled over several years or a single rate for a given time period. In fact, only 32% and 43% of wild and hatchery studies documented estimates on individual years and 8%-10% presented both annual estimates and pooled estimates over a given time period. In total, only 33% of studies provided estimates over several years from the same system (with a maximum of 24 years, Jonsson et al., 2003), allowing appreciation of temporal dynamics and variability of dispersal. These studies suggest strong variation in dispersal rates across years (e.g., Labelle, 1992; Jonsson et al., 2003; Keefer et al., 2005; Fraser et al., 2007; Hess and Matala, 2014), highlighting that a single measure is only a snapshot and should be interpreted cautiously. Multi-year studies open the door to an understanding of drivers of dispersal rates, and how these vary across years with different environmental conditions.

### Implications of dispersal for the conservation and management of anadromous salmonids

Dispersal of anadromous salmonids has implications for both wild- and hatchery-origin fish, as well as metapopulations that include a mixture of wild and hatchery supplemented populations. Our review revealed that most studies discussing the implications of wild-origin dispersal in wild populations focused on one of three primary aspects: demographic, genetic, and/or conservation implications. In terms of demography, the reviewed papers highlighted that dispersal has implications for persistence, stability, and resilience of populations and population complexes. The genetic implications of dispersal that were highlighted largely focused on genetic diversity, and how gene flow alters patterns of genetic diversity across populations, as well as the role of gene flow in constraining or promoting local adaptation. The conservation implications of dispersal tended to focus on the importance of dispersal for recolonization after local extirpation and the potential for population recovery following disturbance (Figure 6). Nevertheless, studies examining the implications of dispersal of wild fish for management of anadromous salmonids continue to be relatively scarce, a point also emphasized by Schtickzelle and Quinn (2007). In fact, only a small proportion of studies focused on dispersal of wild salmonids discussed this topic (26%), and even fewer explicitly evaluated (10%, 15 modelling studies over the 8 species) the implications of dispersal.

**Figure 6:**
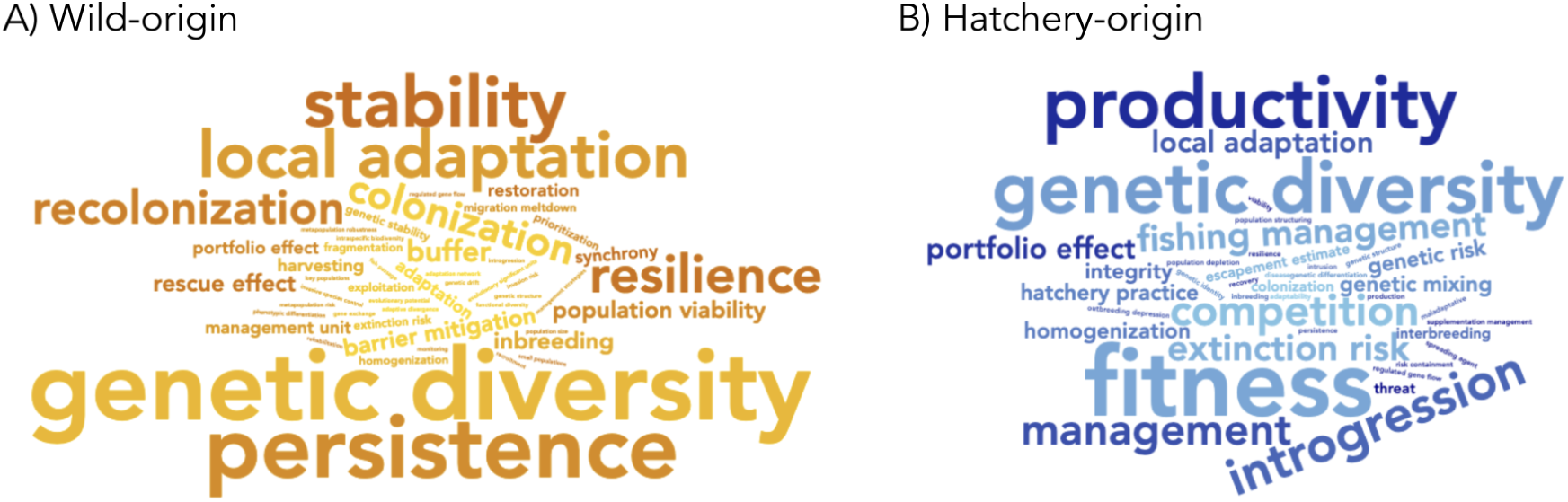
Wordcloud of the different implications of A) wild- and B) hatchery-origin dispersal for conservation and management that were discussed or evaluated by studies. Words size is proportional to their occurrence among empirical and modelling studies.

Yet, neglecting dispersal in metapopulations can affect our understanding of the focal population and the measurement of key demographic parameters used to characterise the status of the population. For instance, return rate is often used as a proxy of survival at sea for anadromous salmonids, which is a crucial indicator of the conditions experienced at sea. It is calculated as the ratio of adults returning in a river and the number of smolts produced for a given cohort. However, return rate is a combination of survival, maturation process, and dispersal. Ignoring dispersal can lead to biased estimates of return rates and survival if the emigration and immigration rates are unbalanced. This, of course, remains a difficult task as it requires monitoring many, if not most, populations and to also then identify the origin of the spawners, both of which are logistically challenging. Dispersal and the functioning of metapopulations also have implications for fisheries management. For example, in the context of source-sink dynamics and metapopulations, source populations can help maintain sink populations that would go extinct in the absence of dispersal. By targeting source populations, which are often the largest, fisheries can affect not only the targeted source populations, but also the sink populations that depend on them. Conversely, any management action that increases or restores a (donor) population may have a positive impact on adjacent (recipient) populations by increasing dispersal. Surprisingly, few studies that we encountered in our review assessed or discussed these potential effects of harvest or management on metapopulation functioning (but see Hindar et al., 2004).

The consequences and implications of hatchery-origin dispersal have received more attention. Indeed, 37% of the hatchery dispersal studies that we reviewed discussed the implications of dispersal. However, only 3% explicitly *evaluated* the implications (and all 3 of these studies used models to assess the implications of dispersal of hatchery fish). Most of the studies discussing implications of dispersal of hatchery fish emphasized consequences for wild populations, highlighting the ongoing tension of conserving wild fish in heavily impacted systems with production hatcheries to support fisheries. Hatchery fish may negatively impact wild populations in a variety of ways, including competition for space or food, attracting predators, as well as concerns about introgression, reduced genetic diversity and effective population size, among others (Naish et al., 2007; McMillan et al., 2023). Several studies included in our review also discuss implications of dispersal of hatchery fish for hatchery management (e.g., adjusting release location and/or timing to reduce straying rates, Lasko et al., 2014; Pascual et al., 1995) and fishery management (e.g., Jonsson et al., 2003). Overall, dispersal of hatchery fish into non supplemented (”wild”) populations complicates the goals of of decreasing fishing pressure on wild (often weaker) populations and increasing it on hatchery (often stronger) populations in hatchery supplemented metapopulations (Knudsen et al., 2021). Hatchery dispersal also make estimates of wild salmon escapement difficult (Keefer et al., 2005; Brenner et al., 2012; Knudsen et al., 2021), and can even mask declines in wild populations (e.g., Johnson et al., 2012). In addition, hatchery broodstock is rarely selected randomly from source populations. Often only anadromous and multiple sea winter fish are used, which are characterised by unique phenotypes and genotypes (e.g., genes associated to age at maturation or migration, Barson et al., 2015; Pearse et al., 2019; Sinclair-Waters et al., 2020), and which favour a particular life history strategy in their progeny. Dispersal of hatchery progeny into wild populations may therefore alter the genetic (e.g., allele frequencies) and life history structure of the recipient populations.

In part because studying dispersal in nature and its effects on salmonid populations is very challenging, modelling approaches have provided a convenient way to simulate sets of interconnected populations and investigate the eco-evolutionary consequences of dispersal, including to evaluate conservation and harvest objectives. We identified 20 modelling studies, most of them focused on Atlantic salmon, sea trout, and Chinook salmon, with a smaller number focused on Coho salmon, Sockeye salmon, and Steelhead trout. A subset of these studies used mathematical and statistical models (e.g., Population Viability Analysis model, graph theory, Stochastic Patch Occupancy Model) to investigate the demographic consequences of dispersal and source-sink dynamics within salmonids metapopulations (Fullerton et al., 2011; Bowlby and Gibson, 2020; Bradford and Braun, 2021). Recently, Lin et al. (2017) and Lamarins et al. (2022) used simulation models (Demo-Genetic Agent Based Model, DG-ABM) of sockeye and Atlantic salmon metapopulations respectively, to evaluate the consequences of dispersal both at demographic, phenotypic and genotypic levels. Castellani et al. (2015) also used a DG-ABM to evaluate the role of immigration on introgression buffering. Metapopulation models have also been developped to evaluate applied management strategies, such as harvesting strategies (Hindar et al., 2004), hatchery intensity (Fullerton et al., 2016), barrier removal (Schick and Lindley, 2007; Ioannidou and O’Hanley, 2019; Rodeles et al., 2021), as well as invasive species regulations (Minett et al., 2023; Healy et al., 2023). Two studies used models to evaluate the fate of wild populations following hatchery fish intrusion (Hindar et al., 2006; Yang et al., 2019). Modelling approaches have been useful in demonstrating the ”dangers” of ignoring metapopulation structure and function (Cooper and Mangel, 1999) and to explore novel management strategies. However, empirical information is ultimately necessary to reproduce a realistic dispersal process within actual metapopulations to help guide appropriate management measures. This body of work highlights the huge promise of teams of empiricists and modelers working together for improving our understanding of dispersal in wild populations and its implications for conservation and management of anadromous salmonids.

## Conclusion

Our review of dispersal in wild- and hatchery origin anadromous salmonids revealed a diversity of studied species, as well as considerable variation in terminology, metrics, and methods for studying dispersal, as well as variation in the spatio-temporal scales considered. Not surprisingly, studies using different approaches (observational, experimental, or modelling), estimation methods, and metrics of dispersal (emigration rate, immigration rate, and gene flow), captured many different facets of dispersal. However, the diversity of approaches and metrics, as well as limited replication for any given method/metric combination, made it difficult to establish a clear relationship between dispersal metrics and drivers across studies. However, the individual studies that have evaluated drivers suggest that different factors matter in different contexts. One of the most consistent results across studies was the influence of transport of hatchery salmon juveniles for release on their tendency to disperse as adults, which has large consequences for wild populations in the same basins (e.g., Sturrock et al., 2019). Overall, our updated review re-confirmed the higher tendency of hatchery-origin fish to disperse than wild-origin fish, however we did find that this pattern did not hold true across all species. However, the unexpected lower immigration rates observed in this review for hatchery-origin fish compared to wild-origin fish require further investigation. We also found that the most robust exploration of the implications of dispersal for conservation and management of salmonids came from modeling studies (e.g., Fullerton et al., 2016; Lin et al., 2017; Ioannidou and O’Hanley, 2019). The integration of empirical knowledge into models and vice versa creates a synergistic relationship that holds considerable promise for highlighting gaps in understanding and motivating further empirical investigation.

There is no single approach or method to fully capture the process of dispersal in wild populations of anadromous salmonids. Instead, it is necessary to combine multiple approaches and to continue to develop (and apply) new methods. For example, recent advances in genetic assignment analysis can be used by employing large numbers of single nucleotide polymorphisms (SNPs) to identify fine-scaled genetic structure across populations and identify immigrants. Additionally, microchemical analysis of calcified structures such as otoliths and ray fins are increasingly used to determine provenance. We also encourage the development (where they do not already exist) of databases of tagging programs from monitored populations that are made publicly available to facilitate the study of wide-ranging salmonids. Advances such as these and others offer exciting opportunities to improve our understanding of dispersal in anadromous salmonids as well as the causes and consequences of dispersal that are difficult to assess directly from a single or a limited number of populations. We recommend that future work continue to explore the influence of dispersal for the dynamics of anadromous salmonid populations and the importance of dispersal as an adaptive life history trait. Furthermore, we encourage the use of generic terms such as dispersal and philopatry rather than straying and homing, although apparent dispersers may sometime be temporary strays. In summary, embracing new methodologies, open data policies, promoting consistency in terminology, and incorporating cutting-edge approaches will be crucial to advancing our understanding of dispersal in wild populations of anadromous salmonids.

## Acknowledgments

We gratefully acknowledge funding from the Region Nouvelle Aquitaine, E2S-UPPA, INRAE and the French Biodiversity Agency (OFB) via the unit Management of Diadromous Fish in their Environment (pôle MIAME). This work was conducted within the International Associated Laboratory MacLife. We thank one anonymous reviewer for comments that greatly improved the manuscript.

## Data availability

The databases used for this review are provided as supplementary materials and are also openly available on a data repository in entrepot.recherche.data.gouv at https://doi.org/10.57745/RCMAGH.

## Funding

We gratefully acknowledge funding from the Region Nouvelle Aquitaine, E2S-UPPA, INRAE and the French Biodiversity Agency (OFB) via the unit Management of Diadromous Fish in their Environment (pôle MIAME).

## Conflict of interest

The authors declare no conflict of interest.

## Appendix S: Supplementary Material

**Figure S1:**
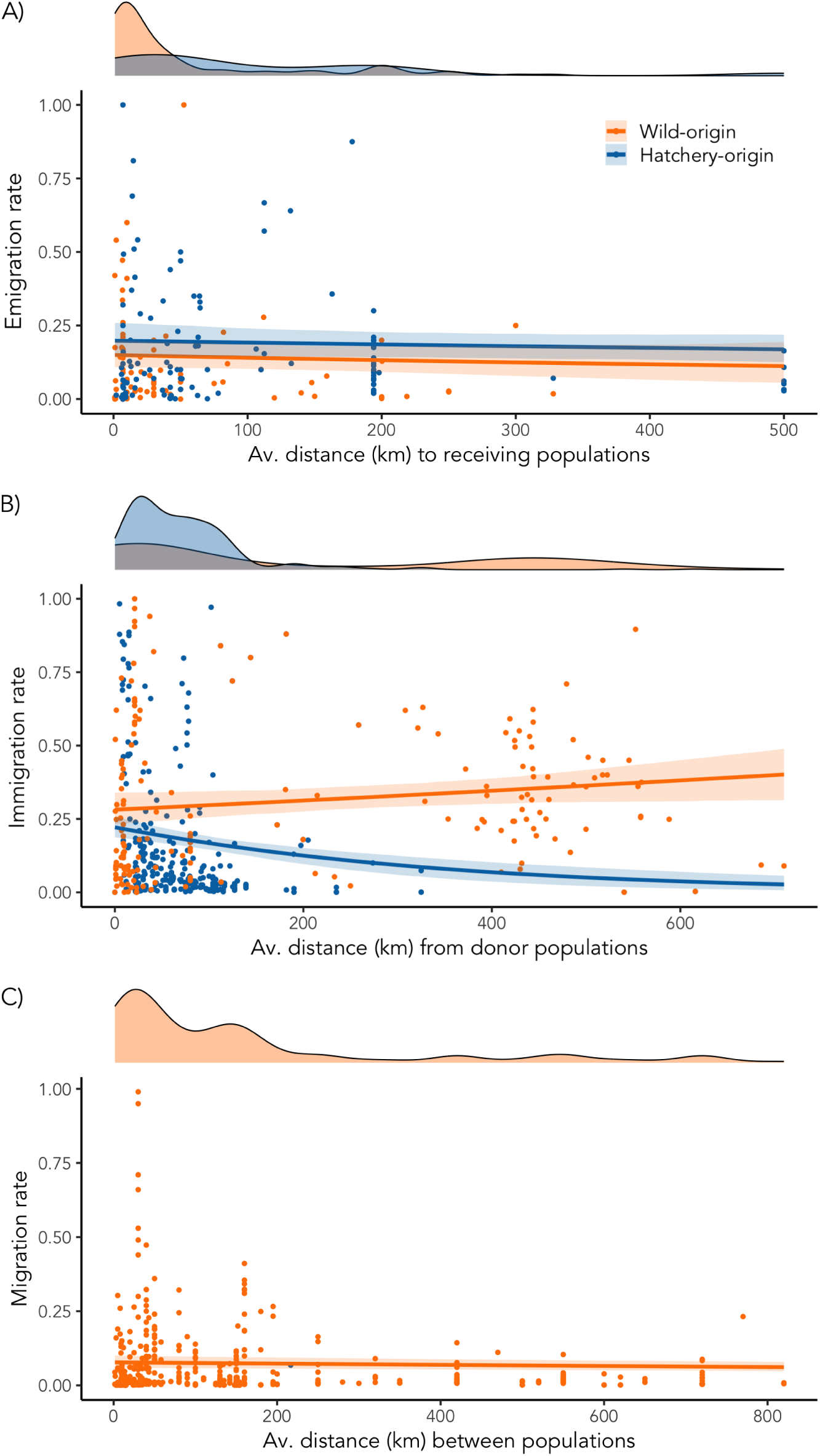
Relationship between estimated A) emigration, B) immigration and C) migration rates over all species and the average distance (km) grouped by the source of fish (wild or hatcheries). Migration rates exclude estimates of number of migrants per generation (*Nm*). Each point is an estimated value. The distribution of distances is also represented in the top of each panel. Fitted curves correspond to beta regressions with logit link (Suppl. Mat Table S2).

**Figure S2:**
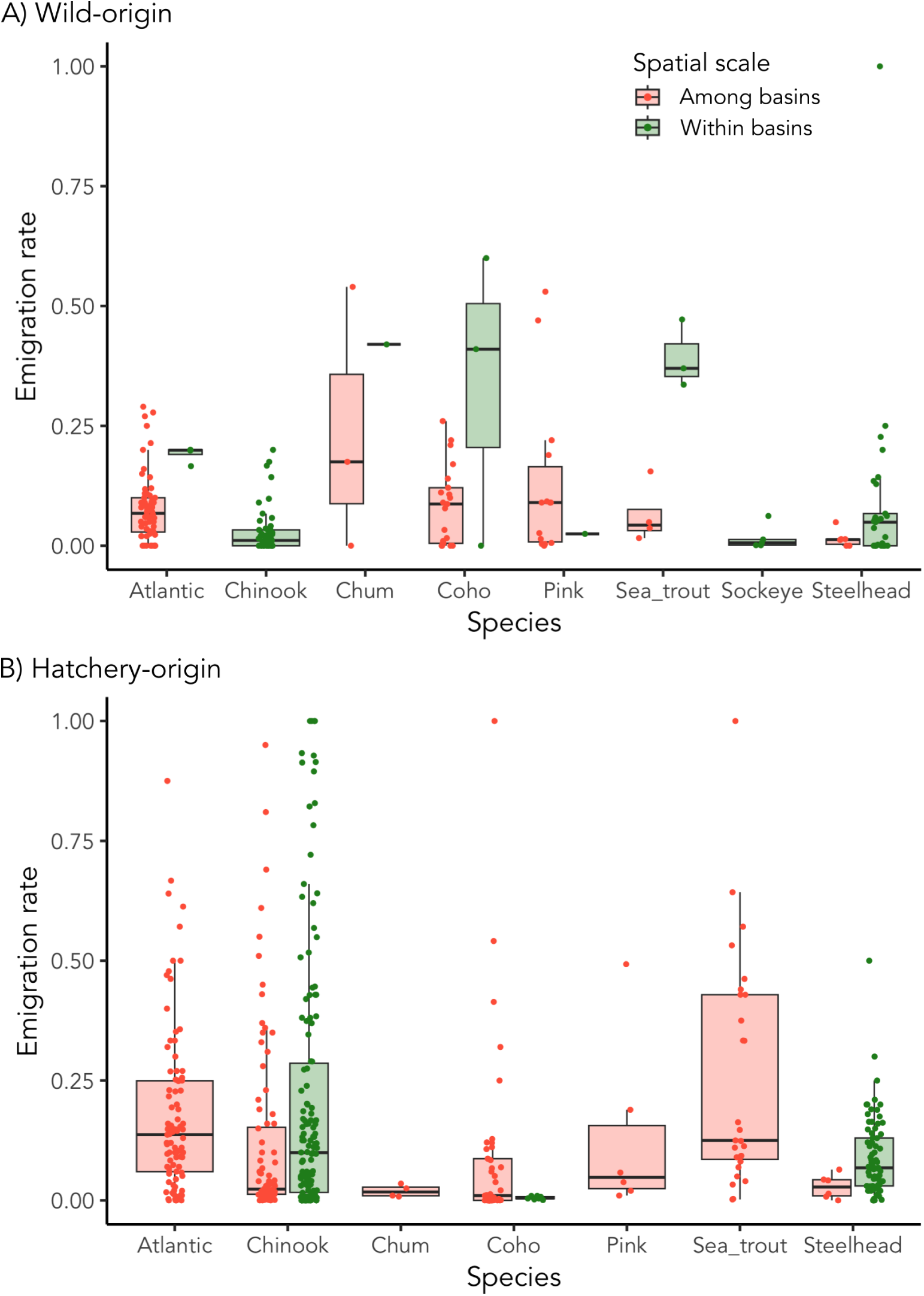
Distributions of estimated emigration rates among species, grouped by the spatial scale of dispersal (among or within basins) for A) wild and B) hatcheries. Each point is an estimated value and the horizontal line shows the median value.

**Table S1:** Similarities and differences between existing reviews. For each study we indicate whether it is a systematic review, the number of species reviewed, whether they differentiate wild *vs.* hatchery fish, whether they reviewed the methods and metrics to estimate dispersal, the factors that might influence its propensity, the estimated rates, but also whether they reviewed modelling studies of dispersal (statistical or simulation models) and implications for conservation. Discuss.= discussed.

| **Review** | Quinn<br>(1993) | Altukhov and<br>Salmenkova<br>(1994) | Hendry<br>(2004) | Schtickzelle<br>and Quinn<br>(2007) | Keefer and<br>Caudill<br>(2014) | Bett et al.<br>(2017) |
| --- | --- | --- | --- | --- | --- | --- |
| **Systematic** | No | No | No | No | No | No |
| **Number species** | 6 | 8 | 8-20 | - | 7 | 7 |
| **Wild *vs* hatchery** | Yes | No | No | No | No | No |
| **Methods** | No | Yes | Yes | No | No | No |
| **Metrics** | No | No | No | No | Yes | No |
| **Factors** | Yes | No | Yes | No | Yes | Yes |
| **Rates estimates** | No | Yes | Yes | No | Yes | No |
| **Models** | No | No | No | No | No | No |
| **Conserv. implic.** | Discuss. | Discuss. | No | Yes | No | Discuss. |

**Table S2:** Summary statistics from the beta regressions with logit link applied to the dispersers proportions, emigration rates, immigration rates, and migration rates.

|  | Estimate | Std. error | z value | p-value |
| --- | --- | --- | --- | --- |
| Dispersers proportion (R2 = 0.1361) |  |  |  |  |
| **Intercept** | -1.051873 | 0.213109 | -4.936 | 7.98e-07 *** |
| **Distance** | -0.003231 | 0.001320 | -2.448 | 0.0144 * |
| **Source** | -0.227149 | 0.261881 | -0.867 | 0.3857 |
| **Interaction** | 0.002306 | 0.001441 | 1.600 | 0.1096 |
| Emigration rate (R2 = 0.04565) |  |  |  |  |
| **Intercept** | -1.7305431 | 0.1385142 | -12.494 | <2e-16 *** |
| **Distance** | -0.0007129 | 0.0013234 | -0.539 | 0.5901 |
| **Source** | 0.3449700 | 0.1753779 | 1.967 | 0.0492 * |
| **Interaction** | 0.0003061 | 0.0015046 | 0.203 | 0.8388 |
| Immigration rate (R2 = 0.1731) |  |  |  |  |
| **Intercept** | -0.9339951 | 0.1062675 | -8.789 | <2e-16 *** |
| **Distance** | 0.0007311 | 0.0003665 | 1.995 | 0.046 * |
| **Source** | -0.3232965 | 0.1366949 | -2.365 | 0.018 * |
| **Interaction** | -0.0040997 | 0.0009464 | -4.332 | 1.48e-05 *** |
| Migration rate (R2 = 0.008781) |  |  |  |  |
| **Intercept** | -2.4719476 | 0.0760418 | -32.508 | <2e-16 *** |
| **Distance** | -0.0003064 | 0.0002301 | -1.332 | 0.183 |

## Notes

### Competing Interest Statement

The authors have declared no competing interest.

### Summary of Updates

A co-author has been added to the manuscript and a comparison between the dispersal from wild- and hatchery-origin salmonids has been implemented, coming with several changes along the manuscript.

